# Spatiotemporal dynamics of calcium transients during embryogenesis of *Drosophila melanogaster*

**DOI:** 10.1101/540070

**Authors:** Olga Markova, Sébastien Senatore, Pierre-François Lenne

**Author notes:** Corresponding authors: OM and PFL.

## Abstract

Calcium signaling plays a crucial role in the physiology of the organs but also in various aspects of the organogenesis of the embryo. High versatility of calcium signaling is encoded by the dynamic variation of intracellular calcium concentration. While the dynamics of calcium is important, little is known about it throughout the embryogenesis of the largest class of animals, insects. Here, we visualize calcium dynamics throughout embryogenesis of *Drosophila* using a fluorescent protein-based calcium indicator, GCaMP3, and report calcium transients in epithelium and neuronal tissues. Local calcium transients of varying duration were detected in the outer epithelium, trachea and neural cells. In addition, gap-junction-dependent calcium waves were identified at stage 16 in the outer epithelium and in the trachea at stage 17. Calcium transient waveform analysis revealed different characteristics as a function of the duration, location and frequency. Detailed characterization of calcium transients during embryogenesis of *Drosophila* will help us better understand the role of calcium signaling in embryogenesis and organogenesis of insects.

## Introduction

The calcium ion is a versatile and universal second messenger involved in the regulation of embryogenesis (Markova et al., 2015, Slusarski and Pelegri 2007, Webb and Miller 2003). Calcium signals have been shown to play a central role in several developmental processes, such as fertilization and organ formation (Kaneuchi et al., 2015, York-Andersen et al., 2015, Christodoulou and Skourides 2015, Berridge, Bootman and Roderick 2003). However, we still lack information about calcium activity during embryogenesis, particularly in non-excitable tissues such as epithelia.

Intercellular calcium waves transmit local information from the initiator cell to a large number of cells in order to coordinate and synchronize their activity during development (Moreno-Juan et al., 2017; Akahoshi, Hotta and Oka 2017, Wallingford et al 2001, Webb and Miller 2003, Chen et al 2017, Wu, Brodskiy, Narciso et al, 2017, Özsu and Monteiro 2017). Surprisingly, endogenous calcium waves during embryogenesis of insects have not been documented. Only local calcium transients were reported during gastrulation (Markova et al., 2015), dorsal closure (Hunter et al., 2014) and during the trachea formation (Caviglia et al., 2016). Here, using GCamp3 as a long-term, calcium-sensitive indicator and confocal imaging, we investigated the dynamics of calcium signaling during *Drosophila* melanogaster embryogenesis. We analyzed calcium transients from the onset of gastrulation to the end of embryogenesis and detected endogenous intercellular calcium waves propagating through gap junctions in the outer epithelium and in the trachea. hese waves occur with a reproducible spatio-temporal pattern, suggesting their potential link to tissue extension, cuticle deposition and neuronal activity.

## Results

### Detected calcium transients

To detect calcium activity in the fruit fly embryo, we ubiquitously expressed GCaMP3 in the embryo using the sqh promoter (sqh::GCaMP3, Markova et al., 2015) and monitored the fluorescence of this indicator throughout embryonic development. Calcium activity is pronounced at particular stages and in specific regions of the embryo. We focused on epithelium tissues and detected the following calcium activity: calcium peaks in the outer epithelium, calcium peaks in the trachea, calcium waves in the outer epithelium, calcium peaks in neurons and calcium waves in the trachea (Figure 1). Below, we characterize each phenomenon separately.

**Fig. 1.**
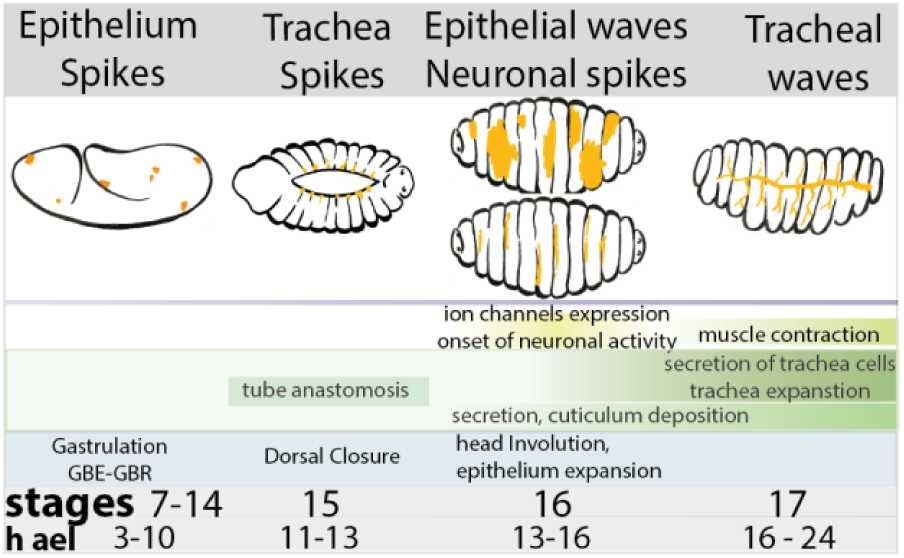
Calcium events during Drosophila embryogenesis

### Calcium spikes in the outer epithelium

To extend the characterization of calcium signals after the stage of gastrulation that we studied previously (Markova et al., 2015) we imaged the outer epithelium until larval hatching and monitored calcium signals at cellular resolution (Figure 1A, SMovie1). We found that each of these transients is spatially restricted to small area of 5-10 µm corresponding to the size of one cell (Figure 2B). Calcium transients were detected in different parts of the outer epithelium thorough embryogenesis (Fig 2C). Temporal projection of calcium spikes shows that calcium spikes are present throughout the outer epithelium (Figure 2D). Spikes last 15 +-7 sec (n = 48 spikes, N = 5 embryos, mean+-SD Fig. 2E). The occurrence of calcium spikes is maximal at gastrulation (3h after egg laying (ael), thereafter the frequency decreases over a few hours before it increases again during dorsal closure (11h ael) (Figure 2F).

**Fig. 2.**
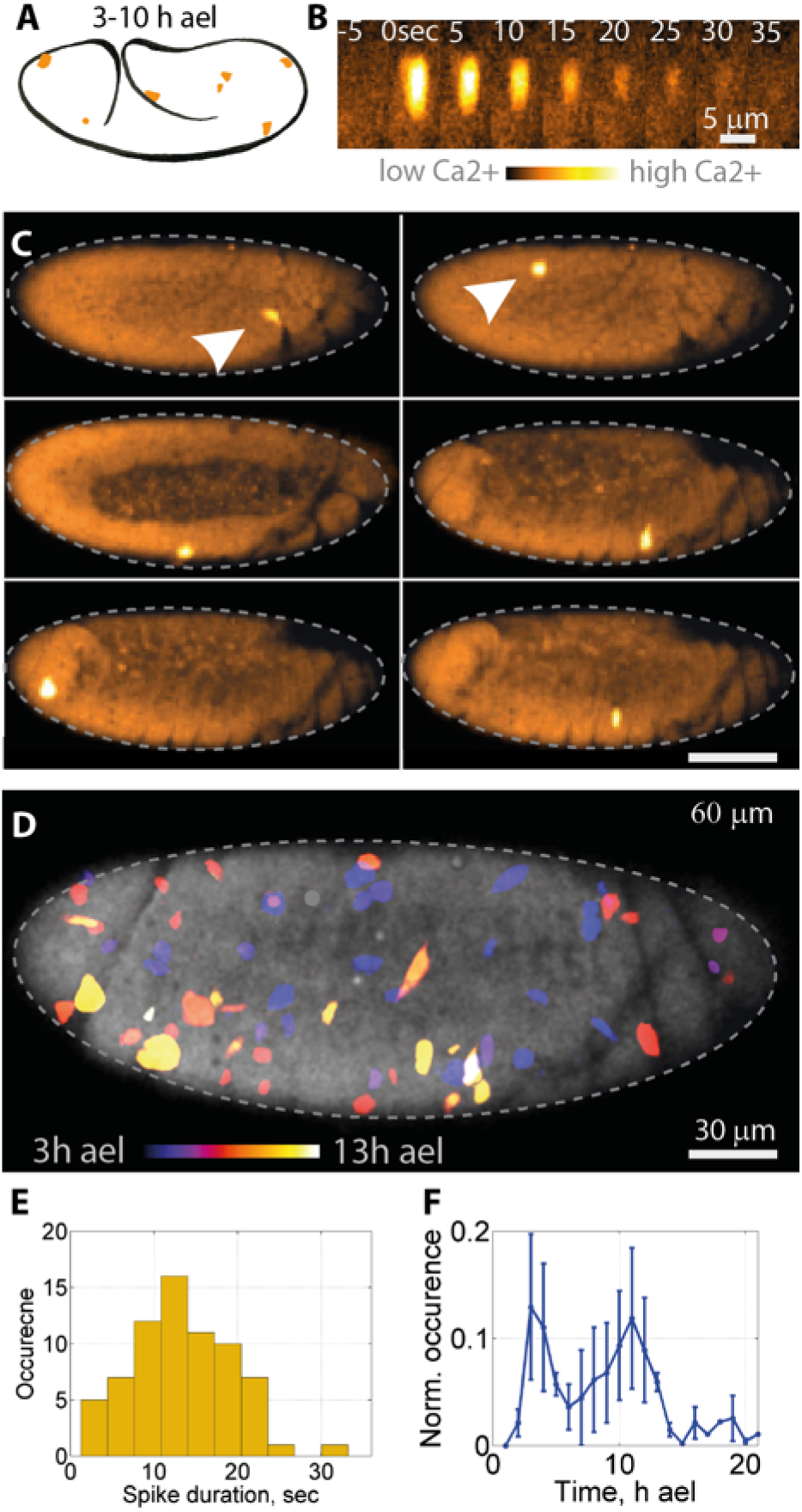
Spikes in the enveloping epithelium. A. Schematic of Drosophila embryo at late gastrulation stage with calcium spikes marked by orange areas. B. Snapshots of a time series showing the appearance and disappearance of one spike. C. Embryo-scale snaphots showing calcium spikes. On the top panels white arrows points to the spikes. D. Color coded projection of all calcium spikes occuring in an embryo during a 10h time-lapse movie. Color codes the time of spikes’appearance. E. Distribution of the duration of calcium spikes. F. Occurrence of spikes during the entire time of embryo development. N = 24 embryos.

### Long calcium spikes

At 9-14 h ael, we detected calcium spikes localized a few micrometers below the embryo surface along the margin between the dorsal ectoderm and the amnioserosa (Figure 3A, SMovie2). By high resolution confocal imaging, we found that many of these spikes arise from cells that have an elongated shape similar to tracheal cells (Figure 3B). While single spikes occurred at different times (Figure 3C, different snapshots), their temporal projection reveals a clear pattern (Fig 2D). Spikes occur on the surface of an ellipse that marks the border between amnioserosa and ectodermal cells (Figure 3D). The duration of these calcium spikes was 120 +/-60 sec (n = 34 spikes) (Fig 3E) that is much longer than those in the ectoderm described above. The location and duration of these long calcium spikes are very similar to those recently reported in the trachea (Caviglia et al. 2016). Spikes occur over a 4-hour period from 9 ael with a maximum frequency at 12 ael (Fig 3F).

**Fig. 3.**
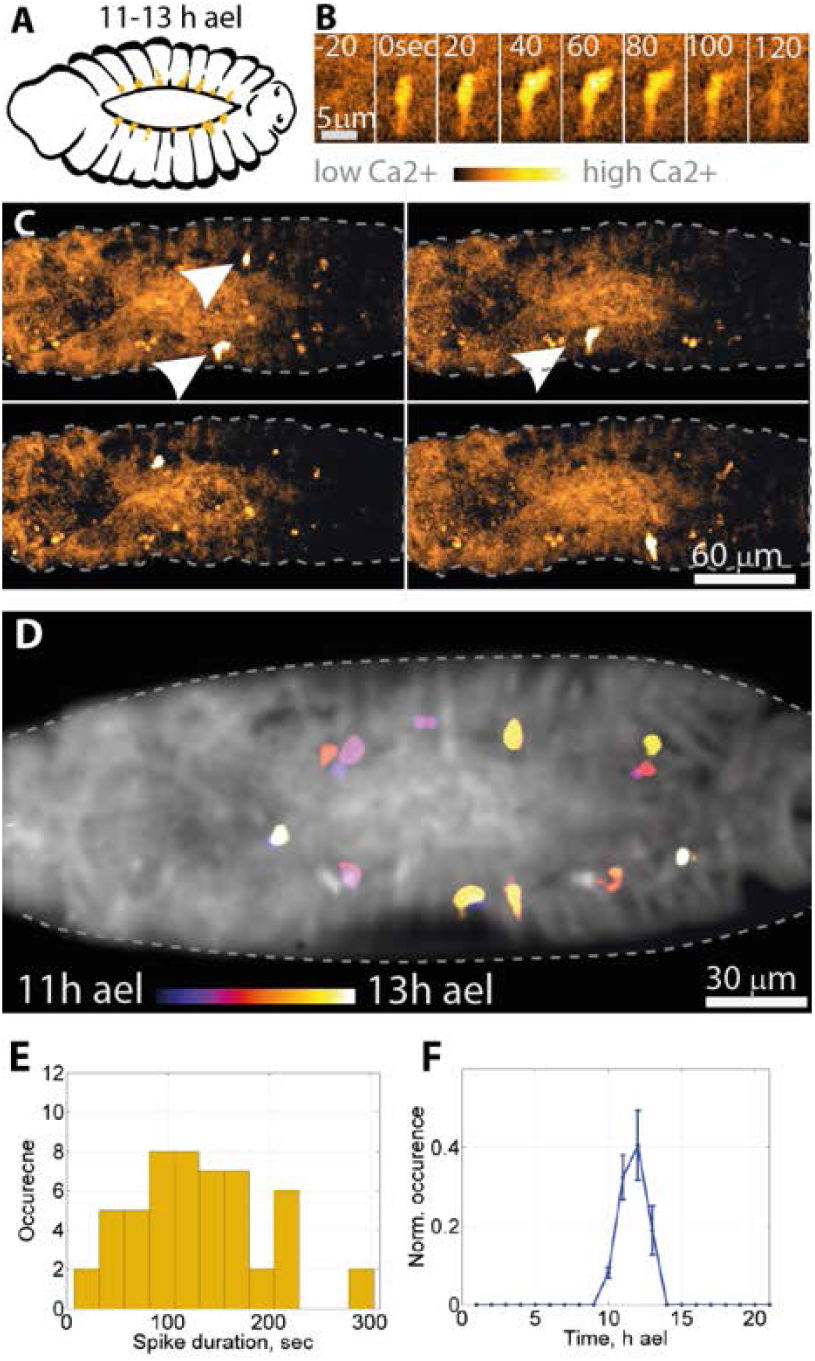
Long calcium spikes. A. Schematic of Drosophila embryo at 11-14 h ael showing the localization of calcium spikes (orange areas). B. Snapshots at 12h ael high resolution of a time series showing the appearance and disappearance of one spike. C. Snapshots of calcium events at the embryo scale. On the top panels white arrows point to the spikes. D. Projection of all calcium spikes occurring in an embryo during a 2h time lapse movie. Color codes the time of spikes’ appearance. E. Distribution of the duration of calcium spikes. F. Occurrence of spikes during embryo development (N = 10 embryos).

### Calcium waves in the outer epithelium

Between 13 and 16 h ael we found calcium waves that propagate in the epithelium at the surface of the embryo (Figure 4A). Calcium waves spread over areas typically 10-30 fold larger than single cell area (5 µm in diameter) (Figure 4B, SMovie3). The propagation velocity of the waves is higher along the dorsal-ventral axis (V_dv = 6.2 +/-3.5 µm/sec, mean+/-SD, n = 115 waves) than along the antero-posterior axis (V_ap = 1.9 +/-0.7 µm/sec, n = 98 waves). Waves propagate in all directions, cross segment boundaries (Figure 4C-D, SMovie4). The duration of the local calcium increase in the one cell area is 14 +/-5 seconds (n = 123 increases, N = 6 embryos, Figure 4E). Calcium waves start at 12h ael, reach a peak of activity at14h ael and fully disappear after 16h ael (Figure 4F).

**Fig. 4.**
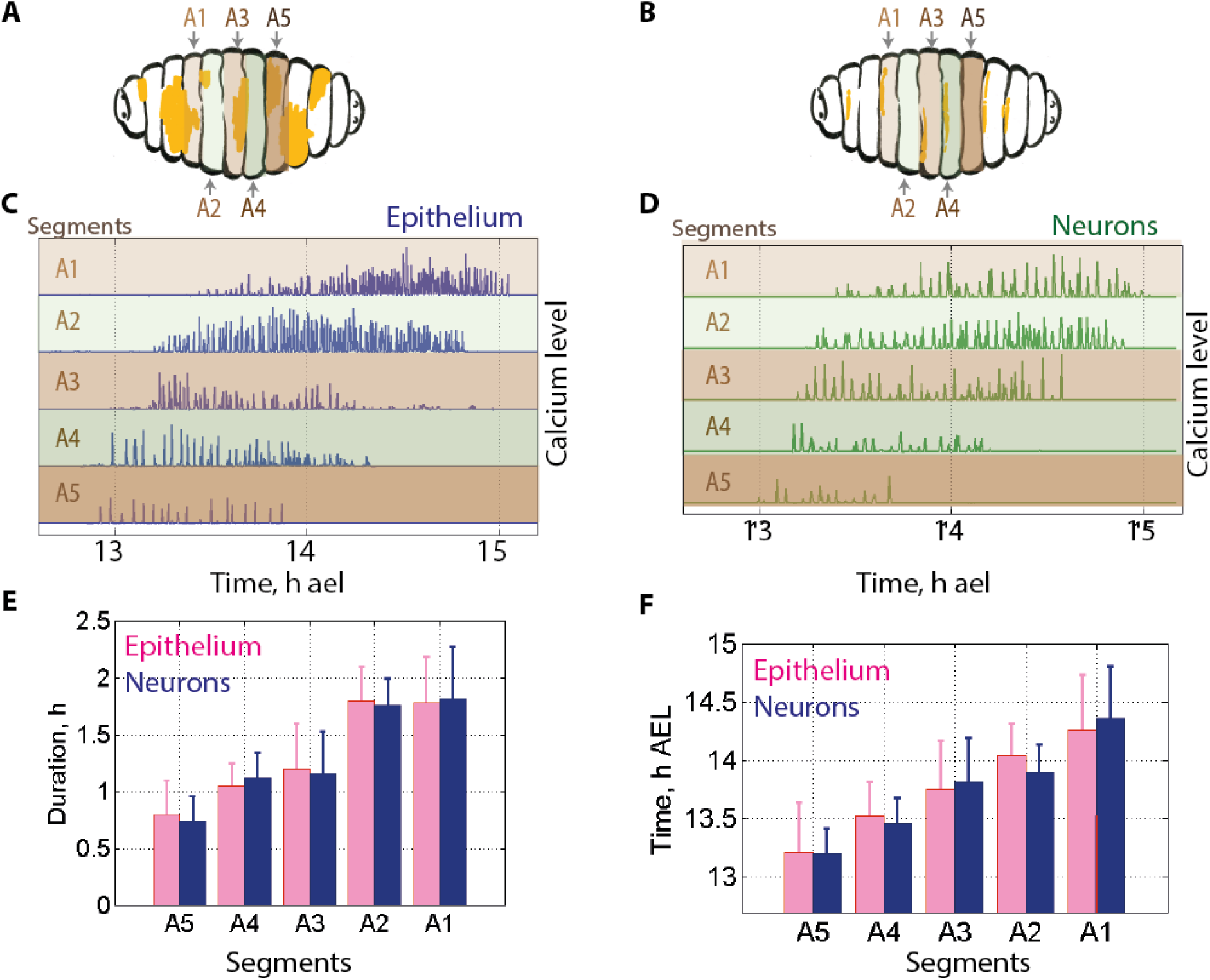
Intercellular calcium waves in the enveloping epithelium. A. Schematic of the Drosophila embryo at 11-16 h ael. Calcium waves are marked by orange color. B. Snapshots of time series showing the propagation of one calcium wave(B) and its color-coded projection over time (B′). C. Snapshots of calcium waves at the embryo scale. D. Color-coded time projection of a 5-minute calcium wave at the embryo scale. E. Distribution of the duration of calcium increases. I. Occurrence of waves during embryo development (N = 14 em bryos). G-1. Calcium activity in innx2 and control embryos. G. Color-coded time projection of a 20-minute recording of the control (69B-GAL4,UAS-GCaMP3) and lnnx2-RNAi embryos (69B-GAL4, UAS-GCaMP3, UAS-lnnx2RNAi). H. Frequency of calcium signalsin WT and lnnx2_RNAi embryos. I. Number of cells particip a­ ting in calcium wave propagation in WT and innx2_RNAi J-K.Localization of calcium wave initiation. J.Kymograph of calcium waves. The localisation of the cell from which the waves started is marked by a white arrowhead. K.Histogram showing the distribution of distances between subsequent origins of calcium waves.

Finally, we noticed that the waves were repeatedly initiated in the same locations (Figure 4J). The initial locations of show high accuracy: they initiated at the same location with variability smaller than a cell size (std = 2.3 µm, N = 95 waves, Figure 4K). These data suggest the presence of an initiator region that generates repetitive calcium waves.

To test if intercellular communication through gap junctions was involved in calcium wave propagation, we knocked-down Innexin which is a major component of gap junctions. Specific knocked down of Innx-2 in the epidermis by RNAi abolished wave propagation, but not transient spikes (Fig 4 G-I). These data indicate that calcium waves propagate in the epithelium via gap junctions.

### Neuronal activity

In addition to the calcium waves that spread on the surface of the embryo, we have detected calcium events located in the neurons underneath the epithelium (Figure 5A). The cells in which calcium events occur have elongated shapes (Figure 5B, SMovie5). Calcium events occur in different segments of the embryo (Figure 5C) and exhibit a regular pattern as shown in the time-projected image (Figure 5D). The duration of the neuronal calcium events is 9+/-7 sec (n = 27 events, N = 4 embryos. Figure 5E). The temporal sequence of these events is as follows: they start at 12 h ael, reach maximum frequency at 14 h ael and disappear at 16 h ael (Figure 5F). This temporal pattern is very similar to that of the epithelium described above.

**Fig. 5.**
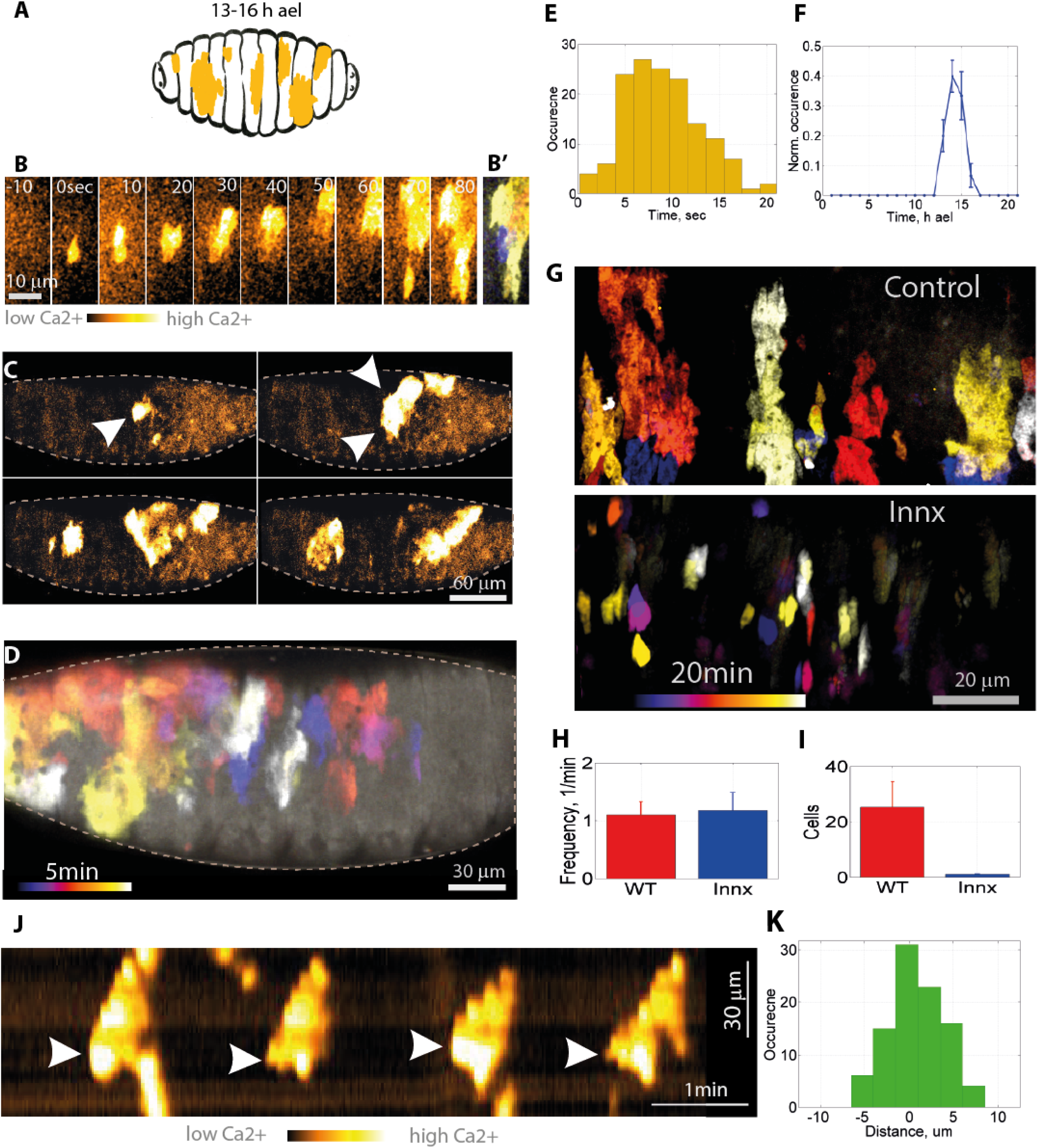
Neuronal calcium activity. A. Schematic of the Drosophila embryo at 11-16 h ael. Calcium signals localised below epithelium are marked by orange color. B. Confocal time series of the calcium activity in the neuronal cells located below the epithelium. C. Snapshots of an embryo producing calcium spikes. D. Color-coded time projection of the neuronal calcium activity at the embryo scale. E. Dura¬tion of calcium spikes. F. Occurrence of spikes during embryo development (N = 7 embryos).

### The spatiotemporal patterns of calcium activity in neurons and in the outer epithelium are not significantly different

Since the calcium waves in the outer epithelium and neuronal calcium activity occur at the same stage of development, we further quantified the extent to which their spatiotemporal patterns are similar. We noted that the spatiotemporal calcium characteristics in neurons and in the outer epithelium depend on the segment in which they occur. We therefore quantified these calcium activities in different segments (Figure 6). Both events occur first in the posterior part of the embryo (Figure 6C, D). They are then visible in the middle and finally in the anterior part of the embryo (Figure 6C, D). The total duration of calcium events varies from 0.75 h in the A5 segment to 1.7 h in the A5 segment (Figure 6E). The development times at which these events occur range from 13.3 h ael in the A5 segment to 14.4 h ael in the A1 segment (Figure 6F). The segment-by-segment comparison of epithelial waves and neuronal activities show no significant difference for all segments (Fig6 C-F). This points to a possible link between these two calcium events.

**Fig. 6.**
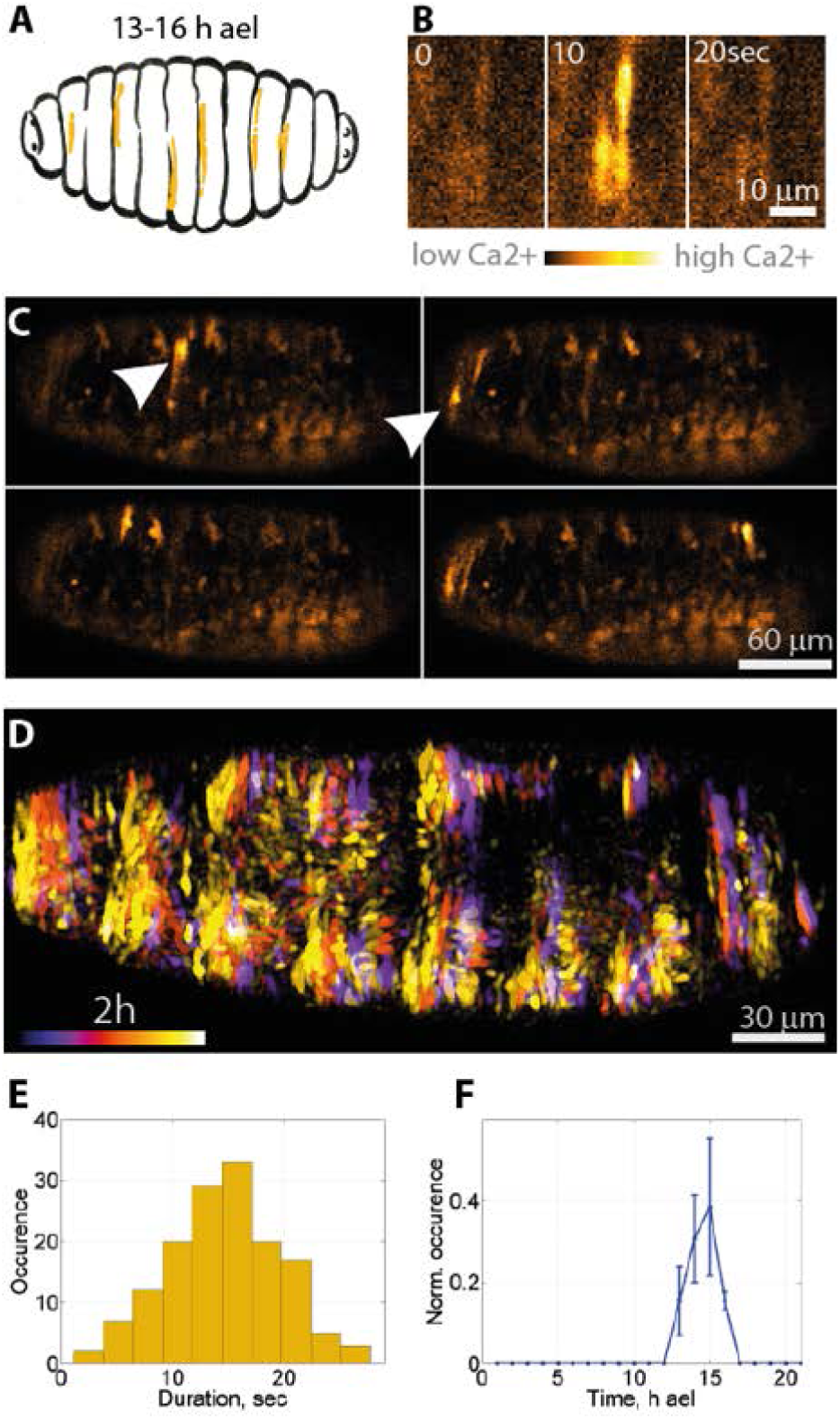
Epithelium waves and neuronal activity. A-В. Schematic of the Drosophila embryo calcium signals at 11 - 16 h ael localised at the embryo surface (A) and 15 um below embryo surface (B). Localization of abdominal segments A1-A5 are pointed by arrows. Calcium events marked by orange color. C-D. Spatio-temporal pattern of epithelium calcium waves (C) and neuronal spikes (D) at segments A1-A5. Graphs show calcium activity for each segment. E. Duration of total period of calcium activity in epithelium and neurons in the segments A1-A5. F. Mean time of calcium activity at segments A1-A5 in epithelium and neuronal cells.

### Tracheal waves

When calcium waves in the outer epithelium become less frequent, the first calcium waves appear in the trachea (Figure 7). The first tracheal calcium waves propagate within the segments and do not pass through the branching points (Figure 7I). A few hours later, calcium waves from the trachea spread along the main trunk and secondary branches (Figure 7B,I, SMovie7). These tracheal waves are produced at intervals with average t = 20 +/-15 min, N = 5 embryos, n = 37 waves, Figure 7I. The temporal projection shows that the waves cover the entire trachea (Figure 7C). Tracheal waves propagate in both directions along the main trunk and secondary branches. The propagation velocity of the calcium wave in the inner segments of the main trunk (V_trunk = 5.2+/-2 µm /sec, N = 51 waves) is lower than that in the secondary branches (V_branches = 9+/-6 µm /sec N = 32 waves). Calcium waves propagating in the main trunk are delayed at each branching point by 6.8+/-2 sec (N = 24 waves).

**Fig. 7.**
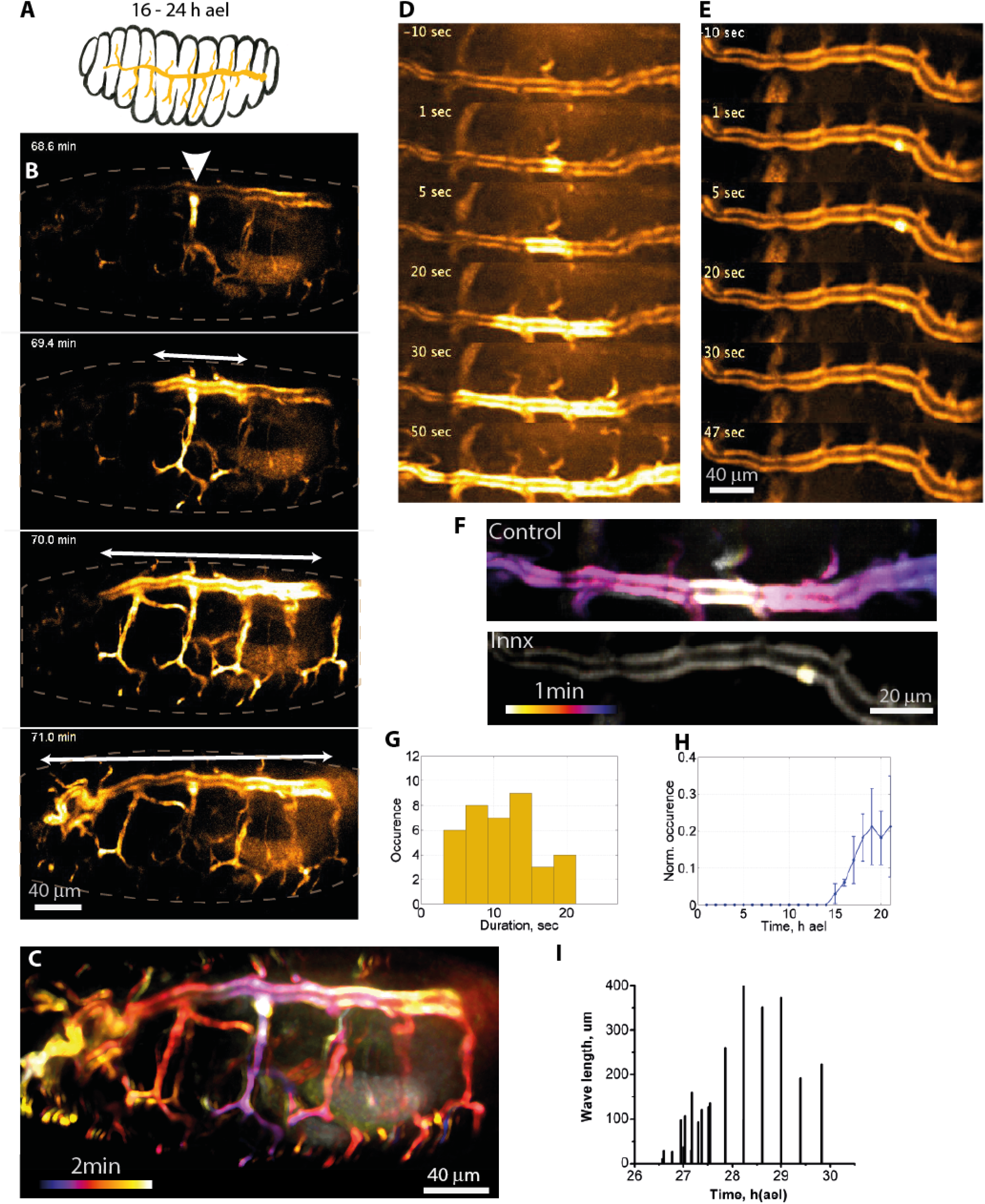
Ca lcium waves in trachea. A. Schematic of the Drosophila embryo at 16-25h ael. Calcium events localised in the trachea are showed by orange color. B. Snapshots of the propagation of the calcium wave in trachea. C. Color-coded tempo­ ral projection of calcium wave propagating along the entire trachea. D-F. Calcium activity in innexin and control embryos. Shapshots of calcium activity in control Brl-GAL4,UAS-GCaMP3(D) and lnnxexin Brl-GAL4,UAS-GCaMP3,UAS-lnnx2RN Ai (F) embryos. F. Color-coded temporal projection of calcium activity in control (Brl-GAL4, UAS-GCaMP3) and lnnx2-RNAi flies (Brl-GAL4,UAS-GCaMP3,UAS-lnnx2RNAi). G. Histogram of durations of calcium increases in trachea cells. H. Occurrence of calcium activity in trachea during embryo development (N = 10 embryos). I. Increase of the duration of calcium wave propagation in trachea towards the end of the embryo development.

To test if intercellular communication through gap junctions was involved in trachea calcium wave propagation, we knocked-down Innx-2 by RNAi in in the trachea, which abrogated wave propagation, but not transient spikes (Figure 7 D-F). These data indicate that calcium waves propagate in the trachea via gap junctions.

The duration of calcium increase within one cell area was 9+/-6 sec (n = 42 events, N = 5 embryos, Figure 7G). Calcium waves in the trachea start at 5h ael and persist until the embryo hatches (Figure 7H).

## Discussion

In this study, we performed long-term calcium imaging of the entire *Drosophila* embryo thorough embryogenesis and characterized the calcium transients from gastrulation to hatching. In addition to the previous local calcium transients observed by the genetically encoded Ca2+ sensor (Markova et al. 2015; Caviglia et al. 2016), calcium activity in neuronal cells, intercellular calcium waves in the epithelium and trachea were identified and characterized by their duration, frequency and distribution (Figure 5).

Epithelial calcium waves that we report here coincide in time with several developmental events: the onset of epithelial secretion and cuticle formation (Bate and Arias 1993), the massive expression of coherent gene expression (Papatsenko, Levine, and Papatsenko 2010) of ion channels producing first neural activity (Baines and Bate 1998) and morphogenic movements that cause head involution (Czerniak et al. 2016). All these events could potentially be related to calcium waves in the covering epithelium.

What determines the time window of calcium waves? The calcium epithelial wave occurs for a short interval at stage 16. It is unlikely that the establishment of gap junctions is involved in this time window, as calcium waves could already be experimentally induced several hours earlier (Hunter et al. 2014). In Drosophila, soluble extract of wing disc triggers calcium epithelial waves (Balaji et al. 2017) and morphogen regulates their amplitude and frequency (Wu, Brodskiy, Huizar, et al. 2017). It is possible that the secretion of a specific molecular component controls the embryonic calcium waves described here. In addition, exogenous release of mechanical load stimulates calcium activity in the embryo (Markova et al. 2015) and the wing disc in the pupa (Narciso et al. 2017). During the period at which epithelial calcium waves occur, the epithelium is actively deformed (Czerniak et al. 2016) and during the tracheal calcium wave period, the muscles strongly contract the embryo. Thus, mechanical stress could potentially activate calcium waves. Finally, the first strong neuronal activity occurs during the epithelial calcium wave period (Baines and Bate 1998). We also show here a striking spatiotemporal correspondence between the neuronal and epithelial activities. The link between these two calcium signals remains to be elucidated.

Calcium waves show distinct speeds of propagation. Epithelial waves propagate more rapidly along the dorso-ventral axis than along the antero-posterior axis. The speed of the tracheal waves also varies as a function of their location in the trachea. It is possible that the differences in wave velocity may be related to differences in cell shape, as reported for wound-induced calcium waves in the wing disk (Narciso et al. 2017).

Although the specific molecular mechanisms underlying calcium signaling associated with *Drosophila* development remain to be elucidated, our quantitative study provides a solid resource to improve our understanding of the role of calcium signaling during insect embryogenesis.

## Methods

### Genetics

Reporter flies (*Sqh::*GCaMP3) were constructed using classic P-element transgenesis as described in (Markova et al. 2015) and used in all experiments except those involving Innx2. The UAS-GCaMP3 fly line was purchased from the Bloomington Drosophila Stock Centre (Fbti0131642); the Gal4-69B fly line was a kind gift of the Razzell Laboratory (Razzell et al. 2013); and the RNAi UAS-dsInnx2 fly line was purchased from the Vienna Drosophila Resource Centre (VDRS Fbgn0027108). Innexin RNAi embryos were obtained by crossing Gal4-69B, UAS-GCaMP3 with UAS-dsInnx2.

### Drosophila care and embryo preparation

The flies were raised at room temperature, while the embryos intended for experiments involving RNAi were raised at 29ºC to increase the activity of Gal4 and RNAi.

### Embryo imaging

Time-lapse movies have been acquired using a confocal spinning disc microscope (Roper) equipped with the following oil-immersion objective lenses (Nikon): 20x 0.75NA Water, 40x 1.2NA Water and 100x 1.4NA Oil. The imaging plan was placed 3-6 μm from the apical surface to image the Calcium signal in the epithelium and 12-20 μm to record the Calcium signal in the trachea and neurons. The temperature during the recordings was maintained at 25°C. The time between frames ranged from 200 ms to 10 s, and the total acquisition time from 1 min to 36 h. The fluorescence of the GCaMP3 gene sensor was excited by a 491 nm laser.

### Image Processing: Temporal Color Coded Projection

Time lapse movies were processed using Fiji as follows: first the Gaussian filter was applied to the whole movie, then the background was subtracted by function Process -> Background Subtraction. Then the difference between subsequent images was calculated using Process -> Image Calculator. Then a mask was applied to identify calcium events. Finally, temporal color image was done using Image -> Hyperstacks -> Temporal Color Code. The latter function attributes a specific color to each frame and project all images into one frame. Color-coded Calcium events were then superimposed on a single frame to show the embryo contours.

### Statistical analysis

Inter-group differences were assessed for significance using the two-sample *t* test and a threshold of *p* < 0.05. Data in graphs are presented as mean ± standard deviation.

### Ethics statement and data availability

This study was carried out in strict accordance with the recommendations in the Guide for the Care and Use of Laboratory Animals of the National Center of Scientific Research (CNRS). No ethics approval was required for this study on *Drosophila*. All data and fly lines described here are available upon request.

## Supporting information

SMovies

SMovies

SMovies

SMovies

SMovies

SMovies

SMovies

## Acknowledgements

This work was supported by an FRM Equipe Grant to P.-F. L. (DEQ201230326509) and an ANR Research grant (‘MORFOR’ Project, ANR11-BSV5-008-01). We thank Dr E. Laugier for technical help. We acknowledge the France-BioImaging/PICsL infrastructure (ANR-10-INBS-04-01) and all members of Lenne’s group. We acknowledge Floris Bosveld, Pierre Mangeol, Jean-Léon Maître and Lena Riabinina for their comments on the manuscript.

## Author Contributions

O.M. designed the project and discussed it with P.-F. L., O.M. performed experiments, analyzed and quantified data, and discussed them with P.-F.L., O. M. wrote the manuscript and P.-F.L. commented on it. S.S. did the sqh::GCaMP fly line and made suggestions for experiments. C.C. quantified data and performed experiments. All authors read and approved the final manuscript.

## References

Akahoshi, Taichi, Kohji Hotta, and Kotaro Oka. 2017. ‘Characterization of Calcium Transients during Early Embryogenesis in Ascidians Ciona Robusta (Ciona Intestinalis Type A) and Ciona Savignyi’. Developmental Biology 431 (2): 205–14. https://doi.org/10.1016/j.ydbio.2017.09.019.

Baines, R. A., and M. Bate. 1998. ‘Electrophysiological Development of Central Neurons in the Drosophila Embryo’. The Journal of Neuroscience: The Official Journal of the Society for Neuroscience 18 (12): 4673–83.

Balaji, Ramya, Christina Bielmeier, Hartmann Harz, Jack Bates, Cornelia Stadler, Alexander Hildebrand, and Anne-Kathrin Classen. 2017. ‘Calcium Spikes, Waves and Oscillations in a Large, Patterned Epithelial Tissue’. Scientific Reports 7 (February): 42786. https://doi.org/10.1038/srep42786.

Bate, Michael, and Alfonso Martínez Arias. 1993. The Development of Drosophila Melanogaster. Cold Spring Harbor Laboratory Press.

Berridge, MJ, MD Bootman, and HL Roderick. 2003. ‘Calcium Signalling: Dynamics, Homeostasis and Remodelling’. NATURE REVIEWS MOLECULAR CELL BIOLOGY 4 (7): 517–29. https://doi.org/10.1038/nrm1155.

Caviglia, Sara, Marko Brankatschk, Elisabeth J. Fischer, Suzanne Eaton, and Stefan Luschnig. 2016. ‘Staccato/Unc-13-4 Controls Secretory Lysosome-Mediated Lumen Fusion during Epithelial Tube Anastomosis’. Nature Cell Biology 18 (7): 727–39. https://doi.org/10.1038/ncb3374.

Chen, Jiakun, Li Xia, Michael R. Bruchas, and Lilianna Solnica-Krezel. 2017. ‘Imaging Early Embryonic Calcium Activity with GCaMP6s Transgenic Zebrafish’. Developmental Biology 430 (2): 385–96. https://doi.org/10.1016/j.ydbio.2017.03.010.

Christodoulou, Neophytos, and Paris A. Skourides. 2015. ‘Cell-Autonomous Ca(2+) Flashes Elicit Pulsed Contractions of an Apical Actin Network to Drive Apical Constriction during Neural Tube Closure’. Cell Reports 13 (10): 2189– 2202. https://doi.org/10.1016/j.celrep.2015.11.017.

Czerniak, Natalia Dorota, Kai Dierkes, Arturo D’Angelo, Julien Colombelli, and Jérôme Solon. 2016. ‘Patterned Contractile Forces Promote Epidermal Spreading and Regulate Segment Positioning during Drosophila Head Involution’. Current Biology 26 (14): 1895–1901. https://doi.org/10.1016/j.cub.2016.05.027.

Hunter, Ginger L., Janice M. Crawford, Julian Z. Genkins, and Daniel P. Kiehart. 2014. ‘Ion Channels Contribute to the Regulation of Cell Sheet Forces during Drosophila Dorsal Closure’. Development (Cambridge, England) 141 (2): 325–34. https://doi.org/10.1242/dev.097097.

Kaneuchi, Taro, Caroline V. Sartain, Satomi Takeo, Vanessa L. Horner, Norene A. Buehner, Toshiro Aigaki, and Mariana F. Wolfner. 2015. ‘Calcium Waves Occur as Drosophila Oocytes Activate’. Proceedings of the National Academy of Sciences of the United States of America 112 (3): 791–96. https://doi.org/10.1073/pnas.1420589112.

Markova, Olga, Sebastien Senatore, Claire Chardes, and Pierre-Francois Lenne. 2015. ‘Calcium Spikes in Epithelium: Study on Drosophila Early Embryos’. Scientific Reports 5 (July): 11379. https://doi.org/10.1038/srep11379.

Moreno-Juan, Verónica, Anton Filipchuk, Noelia Antón-Bolaños, Cecilia Mezzera, Henrik Gezelius, Belen Andrés, Luis Rodríguez-Malmierca, et al. 2017. ‘Prenatal Thalamic Waves Regulate Cortical Area Size Prior to Sensory Processing’. Nature Communications 8 (February): 14172. https://doi.org/10.1038/ncomms14172.

Narciso, Cody E., Nicholas M. Contento, Thomas J. Storey, David J. Hoelzle, and Jeremiah J. Zartman. 2017. ‘Release of Applied Mechanical Loading Stimulates Intercellular Calcium Waves in Drosophila Wing Discs’. Biophysical Journal 113 (2): 491–501. https://doi.org/10.1016/j.bpj.2017.05.051.

Özsu, Nesibe, and Antónia Monteiro. 2017. ‘Wound Healing, Calcium Signaling, and Other Novel Pathways Are Associated with the Formation of Butterfly Eyespots’. BMC Genomics 18 (October): 788. https://doi.org/10.1186/s12864-017-4175-7.

Papatsenko, Ilya, Mike Levine, and Dmitri Papatsenko. 2010. ‘Temporal Waves of Coherent Gene Expression during Drosophila Embryogenesis’. Bioinformatics 26 (21): 2731–36. https://doi.org/10.1093/bioinformatics/btq513.

Razzell, William, Iwan Robert Evans, Paul Martin, and Will Wood. 2013. ‘Calcium Flashes Orchestrate the Wound Inflammatory Response through DUOX Activation and Hydrogen Peroxide Release’. Current Biology: CB, February. https://doi.org/10.1016/j.cub.2013.01.058.

Slusarski, DC, and F Pelegri. 2007. ‘Calcium Signaling in Vertebrate Embryonic Patterning and Morphogenesis’. DEVELOPMENTAL BIOLOGY 307 (1): 1–13. https://doi.org/10.1016/j.ydbio.2007.04.043.

Wallingford, J B, A J Ewald, R M Harland, and S E Fraser. 2001. ‘Calcium Signaling during Convergent Extension in Xenopus’. Current Biology: CB 11 (9): 652–61.

Webb, SE, and AL Miller. 2003. ‘Calcium Signalling during Embryonic Development’. NATURE REVIEWS MOLECULAR CELL BIOLOGY 4 (7): 539–51.

Wu, Qinfeng, Pavel Aleksandrovich Brodskiy, Francisco Javier Huizar, Jamison John Jangula, Cody Narciso, Megan Kathleen Levis, Teresa Brito-Robinson, and Jeremiah J. Zartman. 2017. ‘In Vivo Relevance of Intercellular Calcium Signaling in Drosophila Wing Development’. BioRxiv, September, 187401. https://doi.org/10.1101/187401.

Wu, Qinfeng, Pavel Brodskiy, Cody Narciso, Megan Levis, Jianxu Chen, Peixian Liang, Ninfamaria Arredondo-Walsh, Danny Z. Chen, and Jeremiah James Zartman. 2017. ‘Intercellular Calcium Waves Are Controlled by Morphogen Signaling during Organ Development’. BioRxiv, March, 104745. https://doi.org/10.1101/104745.

York-Andersen, Anna H., Richard M. Parton, Catherine J. Bi, Claire L. Bromley, Ilan Davis, and Timothy T. Weil. 2015. ‘A Single and Rapid Calcium Wave at Egg Activation in Drosophila’. Biology Open, February, BIO201411296. https://doi.org/10.1242/bio.201411296.

