## Supplementary material for "Spatiotemporal dynamics of calcium transients during embryogenesis of *Drosophila melanogaster*": SMovies

### Supplementary Movies

**SMovie1. Spikes in the enveloping epithelium.** 10h time-lapse movie of *Drosophila* GCaMP3::GFP embryo from a gastrulation (3h ael) to dorsal closure (13h ael). 20x Objective.

**SMovie2. Long calcium spikes at tracheal cells.** Time-lapse movie of *Drosophila* GCaMP3::GFP embryo at 11h ael. Dorsal view, 20x Objective.

**SMovie3. Intercellular calcium waves in the enveloping epithelium at the cellular scale.** Time-lapse movie of *Drosophila* GCaMP3::GFP embryo at 14h ael. 100x Objective.

**SMovie4. Intercellular calcium waves in the enveloping epithelium at embryo scale.** Time-lapse movie of *Drosophila* embryo at 14h ael. 20x Objective.

**SMovie5. Neuronal calcium activity.** Time-lapse movie of *Drosophila* embryo at 14h ael. The plane of registration localised below covering epithelium. 20x Objective.

**SMovie6. Calcium waves in trachea.** Calcium wave propagates along the entire trachea. Time-lapse movie of *Drosophila* embryo at 18h ael. 20x Objective.
